# High-resolution image-projection fluorescence lifetime imaging microscopy

**DOI:** 10.64898/2026.06.11.731767

**Authors:** WoongJae Baek, Jongchan Park, Liang Gao

**Affiliations:** Department of Bioengineering, University of California, Los Angeles, California 90095, USA; California NanoSystems Institute, University of California, Los Angeles, California 90095, USA

## Abstract

Fluorescence lifetime imaging microscopy (FLIM) provides molecular contrast that is largely independent of fluorophore concentration, yet it remains constrained by a persistent trade-off among acquisition speed, photon dose, and detector complexity. To address this challenge, we developed image-projection fluorescence lifetime imaging microscopy (IP-FLIM), an integrated optical and computational platform that enables high-resolution, component-resolved lifetime imaging using only a linear single-photon avalanche diode array. We validate IP-FLIM using fluorescent microbeads and bovine pulmonary artery endothelial cells, demonstrating up to 22.3× improvement in contrast-to-noise ratio and 72.3% reduction in background noise over conventional filtered back-projection reconstruction. By combining wide-field projection acquisition with computational *k*-space reconstruction, IP-FLIM provides a scalable route to fast, high-resolution multiplex lifetime imaging.

## Introduction

Fluorescence lifetime imaging microscopy (FLIM) measures the decay rate of fluorophore excited states and provides molecular contrast that is largely independent of fluorophore concentration, excitation intensity, and detection efficiency^1,2,3^. It has been widely applied to autofluorescence imaging of endogenous metabolic cofactors such as NAD(P)H and FAD^4,5^, cancer diagnosis and therapy monitoring^6,7^, and quantitative assessment of protein–protein interactions through Förster resonance energy transfer^2,8^. Moreover, lifetime contrast enables multiplex imaging of fluorophores with overlapping emission spectra but distinct lifetimes, expanding the number of molecular species that can be simultaneously imaged^9,10^.

FLIM techniques are generally classified into two categories: frequency-domain FLIM and time-domain FLIM^11,12^. Unlike frequency-domain FLIM, which measures the fluorescence response at selected modulation frequencies, time-domain FLIM directly records the temporal decay profile following pulsed excitation. This provides richer information about fluorescence decay dynamics and is particularly useful for resolving complex fluorophores or biological autofluorescence signals that exhibit multi-exponential decay behavior.

Conventional time-domain FLIM often relies on point- or line-scanning combined with time-correlated single-photon counting (TCSPC) to acquire lifetime images^1,13^. Although these systems offer high lifetime accuracy at each measured pixel, scanning architectures face two intrinsic limitations. First, each spatial position must accumulate hundreds to thousands of photons for reliable lifetime estimation, which increases pixel dwell time and leads to prolonged acquisition when imaging large fields of view (FOV) or three-dimensional (3D) volumes. Second, the focused laser beams used in scanning approaches can cause photobleaching and phototoxicity, which is particularly problematic for live-cell imaging^14,15^. These constraints have motivated a sustained effort to parallelize FLIM acquisition.

Wide-field FLIM addresses the scanning bottleneck by replacing point detectors with two-dimensional (2D) time-resolved detector arrays that acquire signals from all pixels in parallel. Single-photon avalanche diode (SPAD) arrays have emerged as the leading technology for this purpose, offering picosecond timing resolution, single-photon sensitivity, and per-pixel time-stamping^16,17^. Wide-field SPAD FLIM has been demonstrated with megapixel sensors^18^, time-gated imagers^19^, and dual-gated architectures for snapshot acquisition^20^, and has been integrated with light-sheet microscopy for fast volumetric imaging^21,22^. Despite these advances, most high-resolution 2D SPAD detector arrays operate primarily in time-gated mode because of the engineering challenges associated with integrating per-pixel TCSPC electronics across large arrays. Therefore, scaling wide-field TCSPC FLIM to large FOVs while maintaining high pixel resolution remains technically challenging.

To address this unmet need, we previously developed light-field tomographic FLIM (LIFT-FLIM)^23^, a computational wide-field imaging platform that uses only a linear SPAD array for 3D lifetime imaging. Compared with 2D SPAD arrays, linear SPAD arrays offer a higher fill factor, lower cost, and reduced fabrication complexity^16^, making TCSPC-based FLIM more accessible to general laboratories. However, because LIFT-FLIM relies on aperture division for light-field acquisition, its optical throughput is limited to less than 10%. In addition, the acquisition of sparse tomographic projections can introduce reconstruction artifacts, limiting image quality.

To overcome these limitations, we present image-projection fluorescence lifetime imaging microscopy (IP-FLIM), a 2D counterpart of LIFT-FLIM that provides full optical throughput and an optimized image-processing pipeline for high-resolution imaging of monolayer samples. Like LIFT-FLIM, IP-FLIM acquires fluorescence lifetime information in the spatial-frequency domain by capturing one-dimensional (1D) en-face projections of the sample at a sparse set of projection angles using a linear SPAD array. However, unlike LIFT-FLIM, which transmits only a sub-aperture for subsequent image rotation and compression, IP-FLIM preserves light from the entire pupil, thereby maximizing optical throughput. Meanwhile, a conventional 2D camera simultaneously records a high-resolution reference image of the same sample, which serves as a structural prior for joint reconstruction.

The sparse angular sampling in IP-FLIM requires a reconstruction algorithm that can leverage structural prior information to achieve high-fidelity image recovery. To address this need, we developed a reference-guided *k*-space optimization framework. The algorithm operates entirely in Fourier space and consists of three main steps: first, per-*k* temporal unmixing recovers complex-valued *k*-space coefficients for each fluorescent species at the measured angles; second, a frequency-domain spectral weighting map is derived from the reference-image power spectrum, preserving structurally significant frequencies and suppressing noise-dominated frequencies; and third, the densified total *k*-space coefficients are reconstructed through joint optimization under data-fidelity, spectral-prior, and smoothness constraints.

A key design feature of this framework is that the reference prior is applied only to the total-intensity sinogram, defined as the sum across all lifetime components, whereas the fractional contribution of each component is recovered solely from temporal unmixing. This decoupled strategy improves spatial fidelity without contaminating lifetime contrast, as the reference image provides structural information but does not contain lifetime-specific information that distinguishes different fluorescent species.

### Optical system

The IP-FLIM system uses wide-field illumination to excite the sample (Fig. 1). Fluorescence emission is split into two channels by a beam splitter: 10% of the light is directed to a reference camera, which records a time-integrated fluorescence image over the camera exposure, while the other 90% of the light is imaged by a 4f system with a motorized Dove prism at the Fourier plane. The Dove prism rotates the image by twice its physical rotation angle. The rotated image is then compressed by a cylindrical lens into a 1D projection and detected by a linear TCSPC-SPAD array (SPAD Lambda, PI imaging; 320 pixels).

**Fig. 1.**
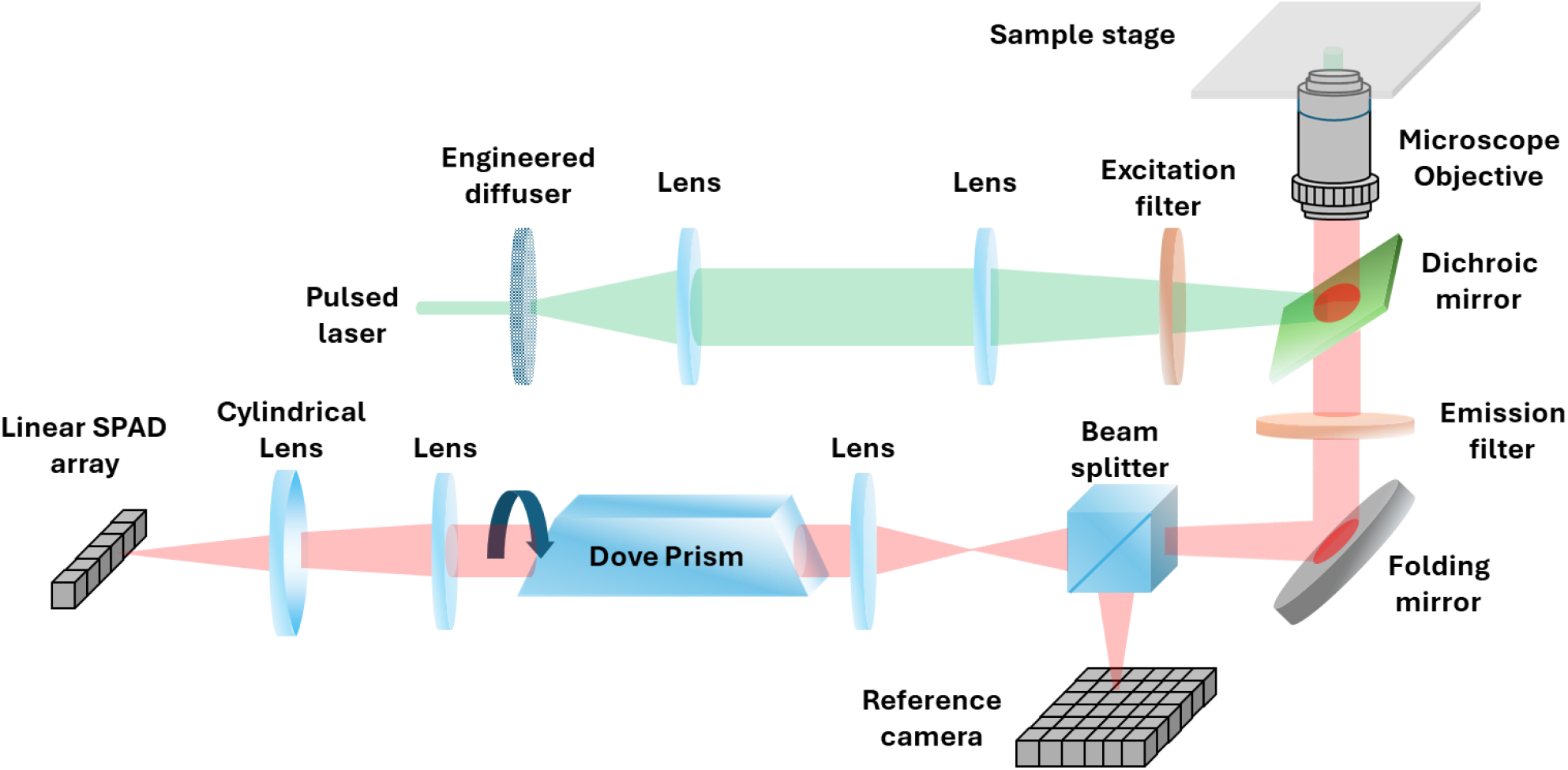
Optical schematic of IP-FLIM. A pulsed supercontinuum laser excites the sample through an inverted epifluorescence microscope. Fluorescence is split by a beam splitter (10:90) between a high-resolution reference camera and a FLIM detection arm. In the FLIM arm, two cascaded 4f relay systems incorporate a Dove prism mounted on a motorized rotation stage to rotate the image, followed by a cylindrical lens that compresses the rotated image into a 1D projection. The projection is detected by a 320-pixel linear SPAD array. Ninety angular projections are acquired sequentially over 0–180° to form the time-resolved sinogram.

We rotate the Dove prism stepwise from 0° to 90° to generate the 0–180° angular span required for tomographic reconstruction. The total acquisition time depends on the number of projection measurements and the exposure time at each angular step. Detailed component specifications, acquisition timing, and IRF measurement are described in Methods.

### *k*-space image fusion for high-resolution image reconstruction

Like sparse-view CT, IP-FLIM acquires image projections from only a subset of angular views, resulting in incomplete sampling in *k*-space. To improve reconstruction quality, we developed a *k*-space image fusion method that uses the high-resolution image acquired by the reference camera as a structural prior.

The reconstruction proceeds in three steps (Fig. 2): per-*k* temporal unmixing, construction of a power spectrum prior from the reference image, and joint *k*-space optimization to produce the component sinograms. The full mathematical formulation, including the forward model, regularization terms, and hyperparameter values, is given in Methods.

**Fig. 2.**
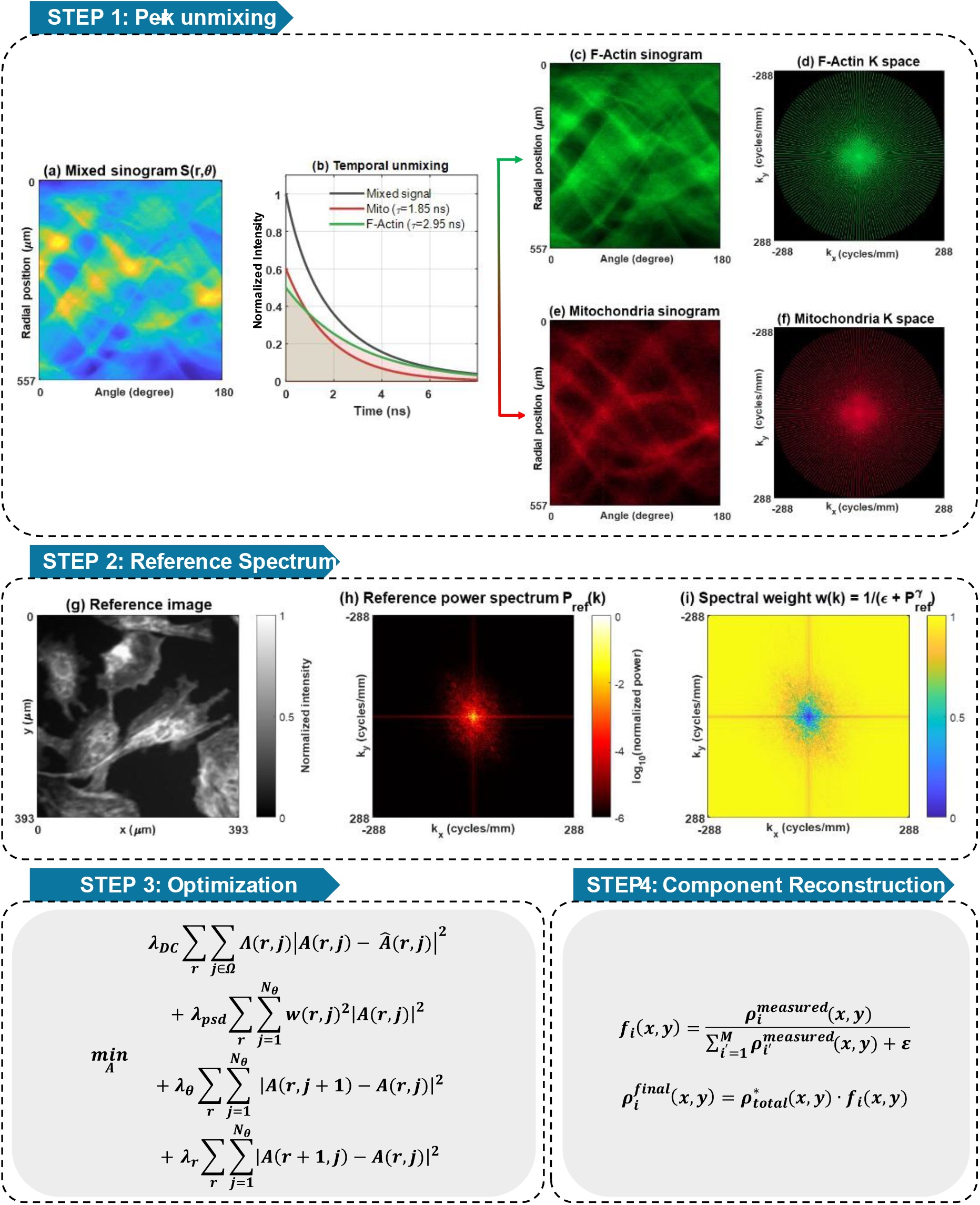
Reference-guided k-space reconstruction pipeline. **Step 1. Per-k unmixing**: (a) measured time-resolved sinogram, (b) temporal dictionary built from the IRF-convolved exponential kernels of the M lifetime species, and the resulting (c,e) per-component sinograms and (d,f) k-space representations. **Step 2. Reference spectrum**: (g) reference image, (h) its 2D power spectrum *P*_*ref*_(k), and (i) the derived spectral weight w(k). **Step 3. Optimization**: total k-space coefficients are jointly optimized on a densified polar grid under data-fidelity (D), spectral-prior (PSD), angular smoothness (θ), and radial smoothness (r) terms. **Step 4. Component reconstruction:** image-domain fraction maps computed from the inverse Radon transform of the per-component sinograms are multiplied with the optimized total image to yield the final component images.

#### Step 1: Per-k temporal unmixing

Let *S(r, θ*_*j*_, *t)* denote the measured time-resolved sinogram, where *r* is the radial coordinate, *θ*_*j*_ is the *j*-th projection angle, and *t* is fluorescence decay time. The signal is modeled as a weighted sum of *M* mono-exponential lifetime components, each multiplied with the corresponding normalized time-resolved fluorescence decay profile. At each spatial location (*r,θ*_*j*_), we solve a regularized linear inverse problem along the time axis to recover the amplitude of each lifetime component, yielding per-component sinograms *S*_*i*_ (*r,θ*_*j*_) for *i* = 1, …, *M* (Fig. 2c, e). Negative amplitudes are clipped to zero to enforce non-negativity. Taking a 1D Fourier transform along *r* then converts each per-component sinogram into its k-space representation (Fig. 2d, f). According to the Fourier slice theorem, each projection angle samples one radial line through the 2D Fourier transform of the underlying component image. This step produces *M* component-specific k-space coefficient maps, each sampled at the original measured projection angles. For comparison, the filtered back-projection (FBP) reconstructions reported below use the same per-pixel temporal unmixing as Step 1; only the subsequent spatial reconstruction differs, ensuring a fair comparison of reconstruction quality.

#### Step 2: Reference power spectral prior

Although the reference image contains no lifetime information, its 2D power spectrum captures the full spatial-frequency content of the sample. We define a power-spectrum weighting function as:

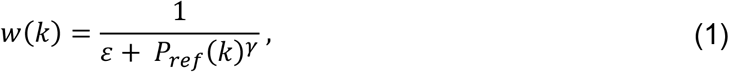

where *P*_*ref*_ (*k*) is the power spectrum of the reference image, and *ε γ* are scalar regularization parameters. This weighting function assigns small values to frequencies with strong structural energy in the reference image, allowing these features to be preserved during reconstruction, while assigning large values to noise-dominated frequencies, which are preferentially suppressed. As a result, the spectral prior adapts automatically to the spatial structure of each sample without requiring frequency-domain tuning (Fig. 2g–i).

#### Step 3: K-space optimization and component reconstruction

We densify the angular sampling and jointly optimize the total *k*-space coefficients *A* (*r,j*) on the polar grid by minimizing an objective function with four terms:

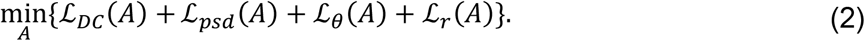

Each term plays a distinct role, as described below.

#### Data fidelity term

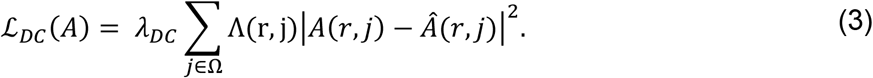

This term constrains *A* (*r,j*) to match the measured *k*-space values *Â* (*r,j*) at the original projection angles indexed by the angular set Ω. The confidence weight Λ (r,j) is derived from the local signal intensity along each measured Fourier slice, so that high-intensity measurements are assigned greater weights than low-intensity measurements.

#### Spectral-prior term

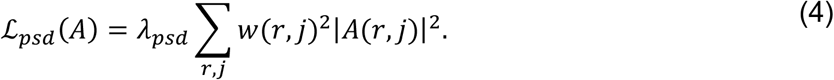

This term suppresses high-amplitude coefficients at noise-dominated frequencies, where the spectral weight *w* is large, while preserving structurally significant frequencies, where *w* is small. Here, *w* is defined on the Cartesian *k*-space grid from Eq. (1) and resampled onto the polar grid. This term transfers the spatial-frequency content of the reference image into the FLIM reconstruction.

#### Angular smoothness term

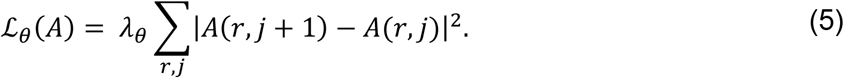

This term promotes consistency between adjacent angular Fourier slices on the densified grid. It propagates information from measured angular slices into the intervening unmeasured regions, thereby filling the angular gaps introduced by sparse-view acquisition.

#### Radial smoothness term

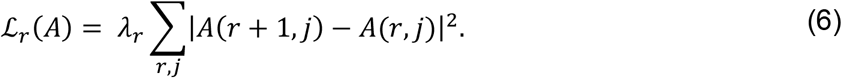

This term suppresses high-frequency noise along the radial direction, providing additional regularization in the radial dimension of polar *k*-space.

#### Solver and densification factor

The four scalar coefficients *λ*_*DC*_,*λ*_*psd*_,*λ*_*θ*_,*λ*_*r*_ control the relative contributions of the four terms. Their values are reported in Supplementary Table S1. These parameters were tuned once using a representative subset of the data and then held fixed across all experiments and simulation conditions reported in this work, so that no per-sample reconfiguration was required.

The full objective function, expressed compactly as Eq. (13) in Methods, is solved by gradient descent with periodic non-negativity projection applied to the inverse-Radon transformed image to enforce physical realizability. The densification factor is a hyperparameter selected to balance angular interpolation accuracy and computational cost. In our implementation, we set the densification factor to eight, as further increases did not appreciably improve the reconstructed images.

#### Step 4: Image-domain fraction and component reconstruction

The optimization in Step 3 is applied only to the total *k*-space coefficients (the sum across all components), reflecting the fact that the reference image observes only the combined fluorescent signal and cannot resolve individual species. To recover individual component images, we adopt an image-domain approach. The per-component sinograms from Step 1 (one per fluorescent species, at the original 90 measured angles) are first inverse-Radon-transformed to image space, yielding measured per-component images 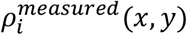. These images carry the species-specific spatial information from the temporal unmixing but are limited by sparse-view artifacts. From them, we compute image-domain fraction maps:

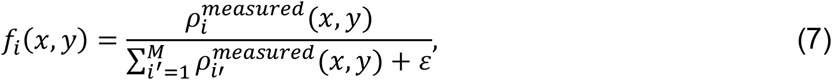

where the sum runs over all *M* species (*i’* = 1, …, *M*) and *ε* is a small constant that prevents division by zero in low-signal regions. The fraction *f*_i_(*x, y*) therefore lies in [0, 1] and represents the relative contribution of species *i* at pixel (*x, y*).

The final per-component image is obtained by multiplying the image-domain fraction with the optimized total intensity image:

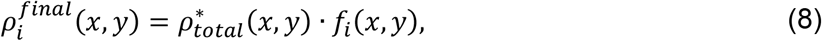

where 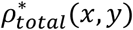 is the inverse Radon transform of the optimized total sinogram from Step 3 (with the asterisk denoting the output of the optimization). The spatial location of each species is determined entirely by the FLIM measurement through temporal unmixing in image space, while the reference image contributes only to the regularization of the total intensity through the *k*-space optimization. This is particularly important for IP-FLIM with sparse projection measurements, where the optimized total-intensity image serves as a weak structural prior to suppress sparse-view artifacts without directly imposing reference-camera structures that may not be supported by the FLIM measurements.

## Results

### Multiplex imaging of fluorescent beads

We validated IP-FLIM on a planar sample containing two species of 15 µm polystyrene microspheres with distinct fluorescence lifetimes (Fig. 3). A total of 90 projections were acquired over an angular range of 0° to 180°. A high-resolution reference image (Fig. 3a) was captured by the reference camera and used both as a structural ground truth and as a spectral prior for image reconstruction. Sample preparation, choice of fluorophore species, and the lifetime values used in the temporal dictionary are described in Methods.

**Fig. 3.**
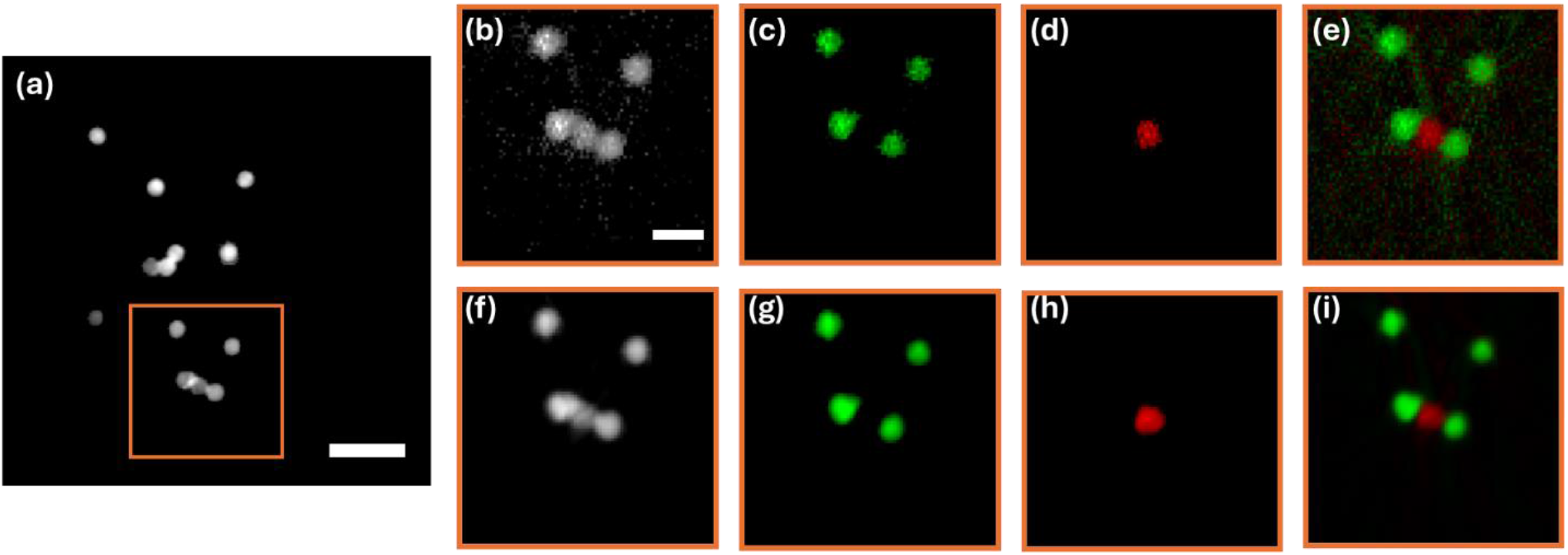
Component separation in fluorescent beads. (a) Reference image acquired by the wide-field reference camera concurrently with the FLIM acquisition. The orange box marks the zoomed-in region in (b–i). (b–e) FBP reconstruction from 90 angular projections: (b) total intensity, (c) Component 1 (yellow-green beads, 505/515 nm), (d) Component 2 (red beads, 580/605 nm), and (e) two-channel composite. (f– i) Corresponding reconstructions by the k-space fusion method: (f) total intensity, (g) Component 1, (h) Component 2, and (i) two-channel composite. Scale bars: 100 µm in (a), 40 µm in (b).

We first reconstructed the component images using standard FBP. Figures 3b–e show the reconstructed total-intensity image, component 1, component 2, and composite image, respectively, within the boxed region in Fig. 3a. Although the bead structures can be recovered, the reconstructions are corrupted by noise and aliasing artifacts arising from the low signal-to-noise ratio (SNR) of the raw TCSPC measurements and insufficient angular sampling.

Next, we reconstructed the images using the *k*-space fusion method (Fig. 3f–i), which substantially improved image quality and reduced reconstruction artifacts. To directly quantify the improvement along structural features, we extracted horizontal line profiles across a region containing closely spaced beads from both species (Fig. 4). The *k*-space fusion method (solid lines) recovers peaks at the correct bead locations with substantially reduced noise compared with FBP (dashed lines). In particular, the FBP reconstruction of component 2 (red) shows large intensity fluctuations across the line profile, whereas the *k*-space fusion method confines the red-channel signal to regions where red beads are physically located. The peak widths recovered by the *k*-space fusion are comparable to those obtained by FBP, indicating that the reference prior does not introduce structures beyond the optical bandwidth of the IP-FLIM system.

**Fig. 4.**
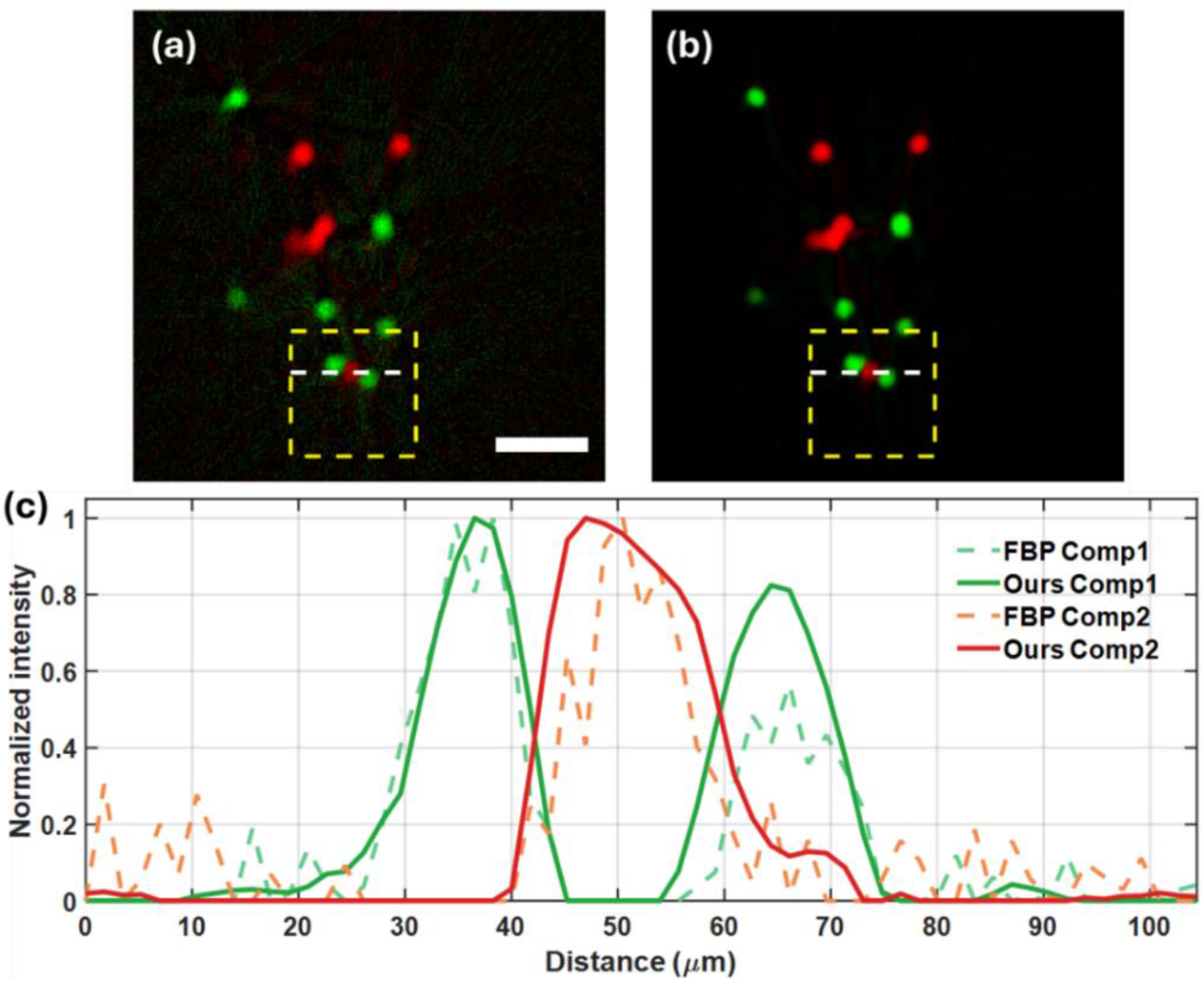
Quantitative comparison of reconstructed line profiles. (a, b) Two-channel composite reconstructions obtained from (a) FBP and (b) k-space fusion. (c) Normalized line profiles of the reconstructed component channels along the white dashed line in (a). Scale bar: 100 µm.

### Multiplex imaging of BPAE cells

To assess performance on biological samples, we imaged bovine pulmonary artery endothelial cells (BPAE) labeled with Alexa Fluor 488 phalloidin (F-actin filaments) and MitoTracker Red CMXRos (mitochondria) (Fig. 5). The IP-FLIM sinogram and the reference image are shown in Fig. 5a and b, respectively.

**Fig. 5.**
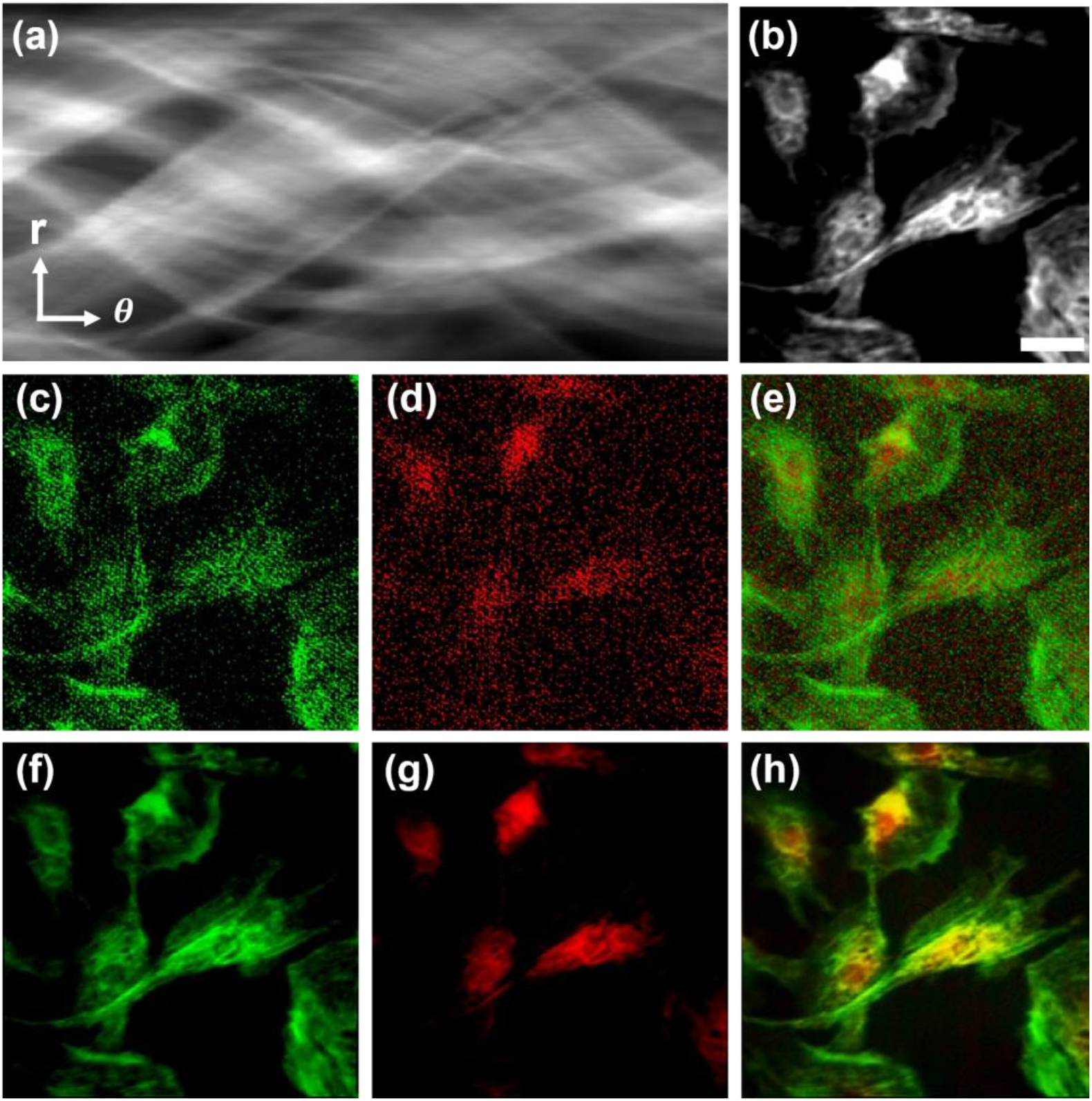
Multiplex imaging of BPAE cells. BPAE cells labeled with Alexa Fluor 488 phalloidin (F-actin filaments) and MitoTracker Red CMXRos (mitochondria), imaged at 40× magnification. (a) Measured time-integrated sinogram. r, radial coordinate; θ, polar angle. (b) Reference intensity image acquired concurrently with the FLIM acquisition. (c–e) FBP reconstruction: (c) F-actin component, (d) mitochondrial component, and (e) two-channel composite. (f–h) Reconstruction by k-space fusion: (f) F-actin component, (g) mitochondrial component, and (h) two-channel composite. Scale bar: 50 µm.

FBP reconstructions of the two component channels are dominated by noise, which obscures the underlying biological structures (Fig. 5c–e). In contrast, the *k*-space fusion reconstruction recovers both fluorescence components with improved image quality (Fig. 5f–g). The composite image (Fig. 5h) shows clear spatial separation between the two species, with green F-actin filaments and red mitochondria resolved as distinct subcellular structures.

Examination of zoomed-in regions from the reference and reconstructed images (Fig. 6) further confirms that the *k*-space fusion method preserves fine structural details. Filamentous structures and mitochondrial puncta visible in the reference images (Fig. 6a,e) are recovered with high fidelity by the *k*-space fusion method (Fig. 6c,g), whereas they are largely unrecognizable in the FBP reconstructions (Fig. 6b,f).

**Fig. 6.**
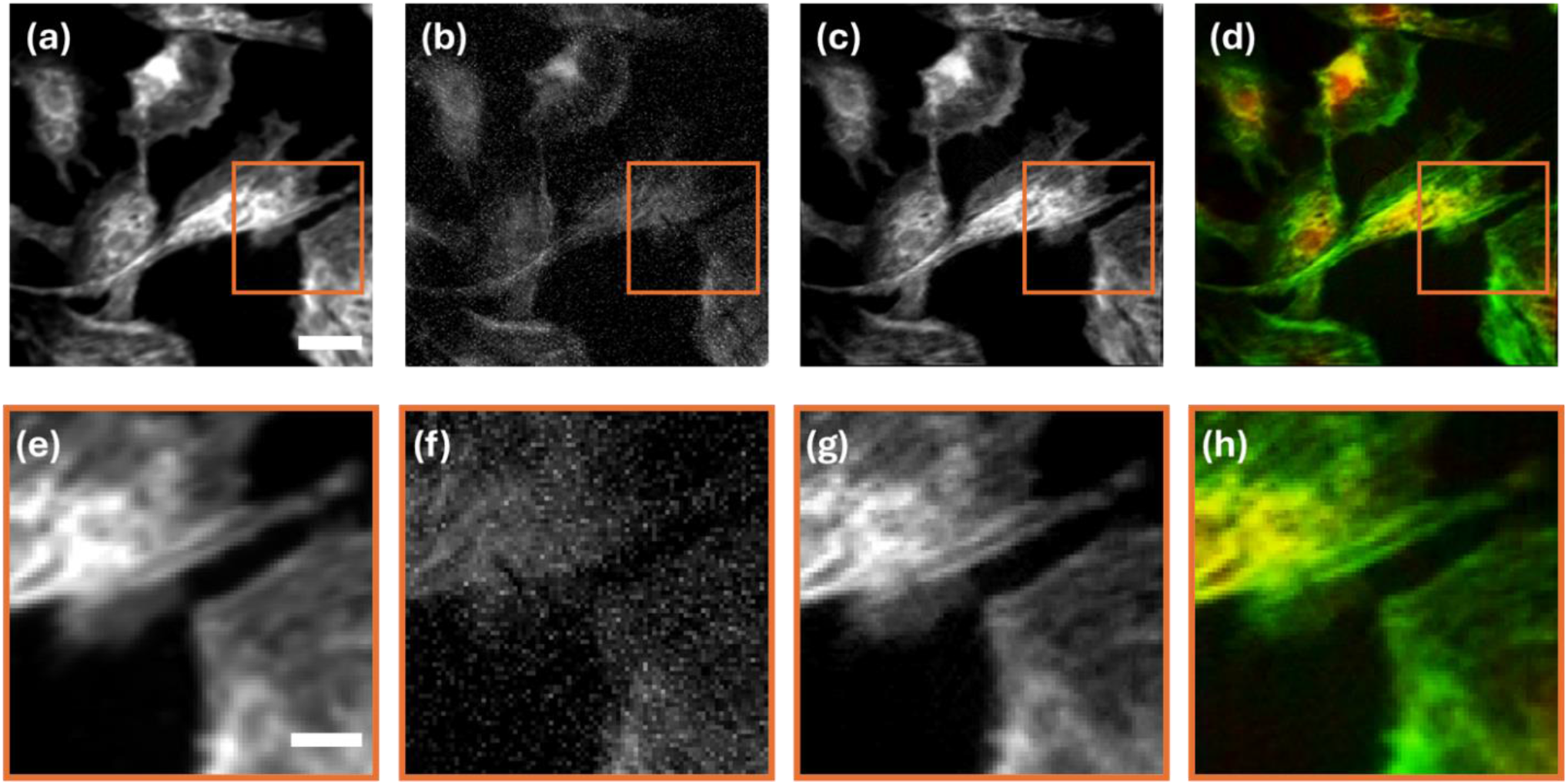
Zoomed-in view of BPAE reconstruction. (a–d) Full-field reconstructions of the BPAE sample: (a) reference image, (b) FBP total intensity, (c) total intensity by the k-space fusion method, and (d) two-channel composite by the k-space fusion method. (e–h) Zoomed-in views of the boxed region: (e) reference image, (f) FBP, (g) the k-space fusion method, and (h) two-channel composite. Scale bars: 50 µm in (a), 20 µm in (e).

### Quantitative reconstruction quality

We quantified reconstruction quality on both samples using four standard metrics: SNR, structural similarity index (SSIM) computed against the reference image, contrast-to-noise ratio (CNR) between the reconstructed signal and background, and the standard deviation of the background intensity (Fig. 7). Definitions, signal and background mask thresholds, and the per-component CNR formulation are reported in Methods.

**Fig. 7.**
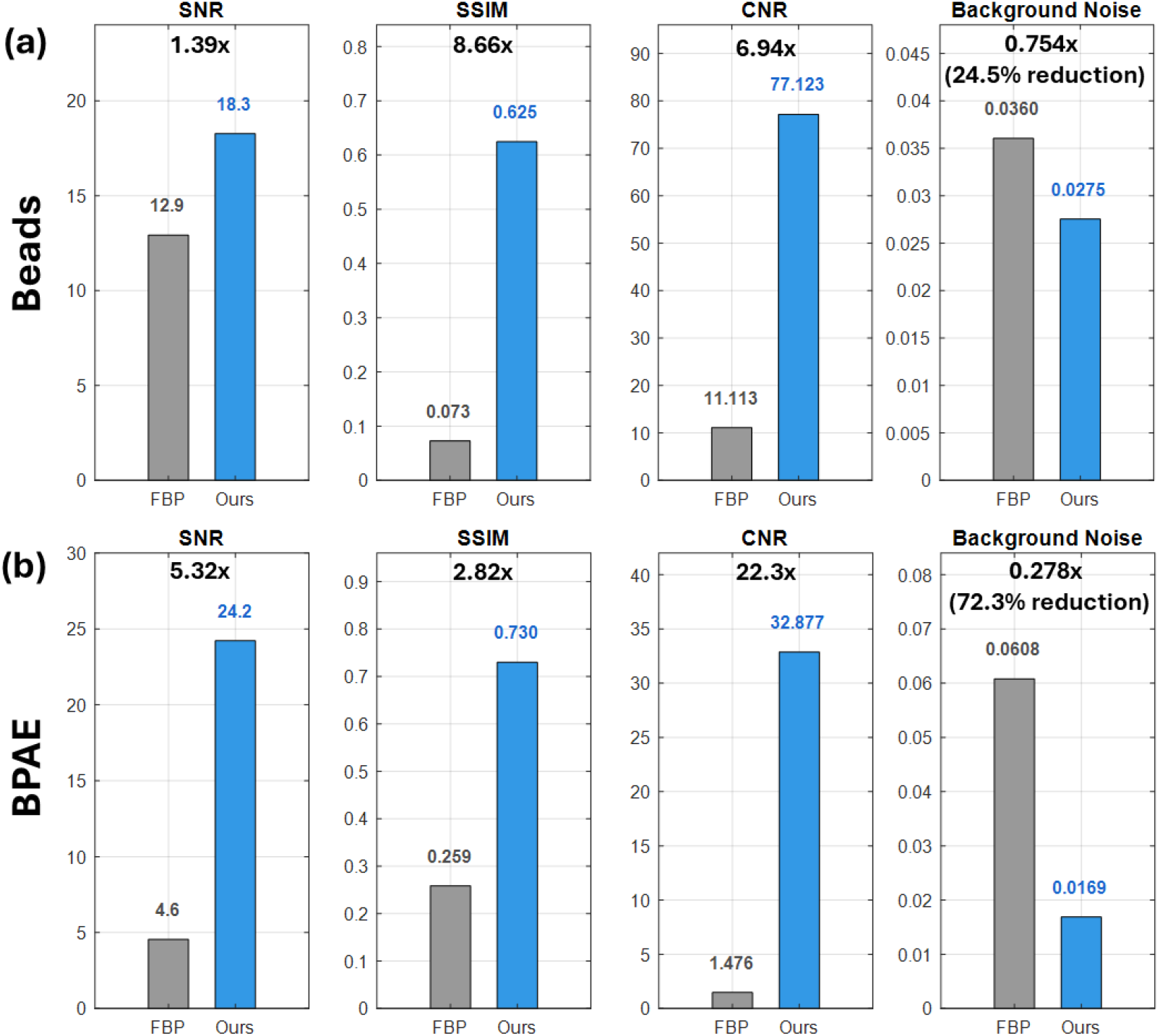
Quantitative comparison of reconstruction quality using standard FBP (gray) and k-space fusion (blue). Signal-to-noise ratio (SNR), structural similarity index (SSIM), contrast-to-noise ratio (CNR), and background noise are compared for (a) fluorescent beads and (b) BPAE cells. Background noise is defined as the standard deviation of pixel intensities in the background region of the normalized reconstructed intensity image.

For the fluorescence bead sample (Fig. 7a), the *k*-space fusion method improves SNR by 1.39×, SSIM by 8.66×, and CNR by 6.94× relative to FBP, while reducing the background-noise standard deviation to 75.4% of the FBP value, corresponding to a 24.6% reduction. The relatively modest SNR improvement reflects the fact that the bead signals are bright and well localized even in the FBP reconstruction. In contrast, the much larger gains in SSIM and CNR indicate that the *k*-space fusion method primarily enhances structural fidelity and background contrast, rather than simply increasing the absolute signal level.

For BPAE cells (Fig. 7b), where the underlying structures are more challenging to resolve, including microscale filaments and puncta against a sample-wide background signal, the improvements are larger across all metrics. The *k*-space fusion method achieves a 5.32× gain in SNR, a 2.82× gain in SSIM, and a 22.3× gain in CNR relative to FBP, while reducing the background-noise standard deviation to 27.8% of the FBP level, corresponding to a 72.2% reduction. These results are consistent with the qualitative observations in Figs. 5–6: under sparse-angle sampling, FBP fails to recover the biological structures because the noise floor overwhelms the filamentous signal, whereas the *k*-space fusion reconstruction transfers spatial-frequency information from the reference image into the lifetime-resolved channels and recovers these structures with high contrast.

The greater improvements observed for the BPAE sample reflect the much poorer FBP baseline in densely structured biological specimens, where sparse-angle streak artifacts overwhelm true cellular structures. In addition, the dense cellular architecture provides richer spatial-frequency content in the reference image, allowing the spectral prior to more effectively suppress artifacts and preserve fine structural features. In contrast, for the sparse bead sample, FBP already recovers bead positions reasonably well because the beads’ *k*-space power spectrum is concentrated near the origin, allowing them to be sampled adequately even with sparse angular projections. This differs from BPAE cell structures, which contain much higher spatial-frequency information and are more susceptible to sparse-view artifacts.

Beyond these two experimental samples, we further characterized the operating regime of the *k*-space fusion method through numerical simulations by varying both the lifetime separation between species and the number of fluorescent species while keeping all other parameters fixed (Supplementary Information). The *k*-space fusion method maintains substantially above-chance per-component contrast across all tested conditions and degrades smoothly as lifetime separation decreases or the number of species increases. These simulations provide a quantitative assessment of the conditions under which the reconstruction is expected to remain reliable.

Across both experimental samples, the same four metrics consistently favor the *k*-space fusion method by factors that depend on the structural complexity of the sample. The benefit was greatest for biological imaging scenarios involving micron-scale, low-contrast structures embedded in heterogeneous backgrounds. These results confirm that reference-guided reconstruction extends the practical operating regime of sparse-view IP-FLIM, enabling reliable imaging of biological samples that would otherwise be inaccessible to FBP-based reconstruction.

## Conclusion

The IP-FLIM platform presented here represents a computational FLIM modality for high-resolution, high-throughput cellular imaging. By combining sparse-view acquisition with *k*-space fusion reconstruction, IP-FLIM achieves both improved imaging throughput and high reconstruction quality. We anticipate that the jointly designed optical front end and reconstruction algorithm, as exemplified by IP-FLIM, will be applicable to a broad range of lifetime imaging applications, enabling component-resolved lifetime contrast over wide fields of view using cost-effective 1D ultrafast detector-array hardware.

Several limitations remain. First, the current implementation assumes a fixed number of lifetime components and known component lifetime values. This condition is well suited for multiplex imaging, where the fluorophores used to label the sample and their approximate lifetimes are known in advance. Moreover, the procedure described in Methods further estimates these lifetime values directly from the measured sinogram, requiring only the number of components as input. Extending IP-FLIM to samples in which the number of underlying lifetime species is not known *a priori* would require a blind unmixing^24,25,26^ step to estimate both the number and the lifetime values of the components from the data. Second, the optimization is currently solved by gradient descent with empirically tuned hyperparameters. Data-driven approaches, such as learned priors or unrolled optimization networks^27-30^, could potentially further reduce the number of measured projections required to achieve a desired reconstruction quality. Finally, IP-FLIM is currently applicable to only planar static samples. Extending the approach to volumetric imaging through axial scanning, and to dynamic imaging using fewer projection angles, represents important directions for future investigation.

## Methods

### Optical system

#### Excitation and microscope front-end

A pulsed supercontinuum laser (SuperK FIANIUM FIU-15, NKT Photonics) operating at a 39 MHz repetition rate serves as the excitation source. Excitation wavelengths were selected by a multi-band excitation filter (ZET405/488/561/647x, Chroma): 488 nm and 561 nm were used simultaneously for BPAE imaging, and 505 nm and 580 nm were used for the fluorescent beads. The 405 nm band of the excitation filter was not used because the supercontinuum source is bandpass-restricted to wavelengths above 460 nm; consequently, DAPI in the BPAE slide was not excited and does not contribute to either the reference image or the FLIM channels. The laser is coupled into an inverted epifluorescence microscope (IX83, Olympus) and focused onto the sample by a 20× objective lens (UPLXAPO20X, Olympus). For BPAE imaging, an additional 2× beam expander was inserted in the detection path, giving an effective magnification of 40× and a sample-plane pixel size of 0.87 µm; for the beads, the original 20× configuration was used, giving a pixel size of 1.74 µm. Excitation is separated from fluorescence by a multi-band dichroic mirror (ZT405/488/561/647rpc, Chroma) and emission is filtered by a multi-band emission filter (ZET405/488/561/647m, Chroma).

#### Detection beam splitting

The intermediate fluorescence image at the side image port of the microscope is split by a non-polarizing plate beam splitter (BSX16, Thorlabs), which transmits 10% of the light to a high-resolution reference camera (CS2100M-USB, Thorlabs; 1920 × 1080 pixels) and reflects 90% of the light to the FLIM detection arm. The asymmetric splitting ratio favors photon collection efficiency in the FLIM channel, where photon counts directly determine the accuracy of lifetime estimation, while the longer integration window of the reference camera (5 s) provides sufficient signal for the structural reference.

#### Spatial encoding via Dove prism and cylindrical lens

The FLIM detection arm performs angular projection acquisition using two cascaded 4f relay systems that propagate the intermediate image from the microscope side port to the linear SPAD array. The first 4f relay consists of the microscope tube lens (*f* = 180 mm, Olympus) and an 80 mm achromatic lens (Thorlabs), forming an intermediate pupil (Fourier) plane at which a Dove prism (PS990M, Thorlabs) mounted on a motorized rotation stage (DDR100, Thorlabs; maximum rotation speed 180 rpm) is positioned. The second 4f relay consists of an 80 mm and a 150 mm achromatic lens (Thorlabs), forming a second image plane downstream of the Dove prism. Rotation of the Dove prism by an angle *φ* rotates the relayed image by 2*φ*, so that a physical rotation range of 0–90° generates image rotations spanning 0–180°, which is the full angular coverage required for tomographic reconstruction. A cylindrical lens (*f* = 19 mm; Thorlabs) with its invariant axis aligned to the linear SPAD array is placed at an appropriate distance after the second 4f relay. The cylindrical lens compresses the rotated image along the axis perpendicular to its invariant axis onto the linear SPAD array, producing a 1D projection at the corresponding rotation angle. The axial positions of the cylindrical lens and the SPAD array were adjusted to minimize the line-image width.

#### Time-resolved detection

The line projection is detected by a linear SPAD array (SPAD Lambda, Pi Imaging Technology) with 320 × 1 active pixels at a 29 µm pitch. Each pixel integrates a TCSPC histogram with 20 ps timing resolution, a peak photon detection probability of 50% at 520 nm, a fill factor exceeding 80%, and a median dark count rate below 250 counts per second per pixel. The maximum photon throughput in time-tagging mode is 140 Mcps. Combined with the 39 MHz laser repetition rate, the detector observes one excitation cycle every 25.6 ns.

### Data acquisition

#### Acquisition timing

For each rotation angle, the SPAD array integrates photons for an exposure time of 100 ms. A 100 ms idle interval was inserted between exposures, dominated by USB data-transfer overhead from the SPAD module to the host computer that must complete before the next exposure begins, with a smaller contribution from rotation-stage settling. A complete dataset consisting of 90 angles uniformly distributed across 0–180° (corresponding to 0–90° physical rotation of the Dove prism) was acquired in approximately 21 s. The reference camera operated concurrently with a 5 s integration time and accumulated the multi-band intensity image of the same field of view over multiple FLIM exposures. The current acquisition speed is limited by this data-transfer overhead rather than by photon counts; faster readout interfaces (such as PCIe-based data streaming in place of USB) or a coarser temporal binning that reduces the per-frame data volume would directly reduce the per-angle idle interval and therefore the total acquisition time, in principle to near-photon-limited operation.

The choice of 90 angular projections is a design parameter rather than a hardware constraint of the system. The Dove-prism rotation stage supports arbitrary angular sampling, and the reconstruction algorithm is agnostic to the specific number of projections. Ninety projections were selected as a practical balance between acquisition time and angular sampling density: this number provides 2° angular spacing across the 0–180° range, which we found empirically sufficient for the reference-guided reconstruction to recover fine biological structures (Figures 5 and 6) while keeping the total acquisition time within a range compatible with potential live-cell imaging extensions. Both fewer projections (for faster acquisition at the cost of reconstruction quality) and more projections (for improved reconstruction at the cost of speed) are readily supported by the same system, and the operating regime of the algorithm as a function of angular sampling density is important to investigate but outside the scope of the current study.

#### Raw data structure and temporal preprocessing

The raw SPAD output is structured as a tensor with dimensions (320 radial positions × 2560 time bins × 90 angles). Each time bin is 10 ps wide, giving a total observation window of 25.6 ns per histogram. To increase photon counts per bin and reduce computational cost, the time axis was rebinned by a factor of 25.6 to 100 bins, each 256 ps wide. A subset of 60 bins (bins 11–70 after rebinning) covering the rising edge and the dominant decay region of the fluorescence signal was retained for reconstruction; bins outside this range contained either pre-pulse noise or low-amplitude tail and were discarded. The resulting input to the reconstruction pipeline is therefore a (320 × 60 × 90) tensor.

#### Instrument response function

The instrument response function (IRF) was characterized by replacing the sample with a planar mirror at the sample plane and recording the back-reflected laser pulse with the same SPAD acquisition settings used for fluorescence imaging. The measured IRF is assumed to be invariant across the 90 rotation angles, since rotation of the Dove prism does not introduce angle-dependent timing distortions in our optical configuration. The IRF was used to construct the temporal dictionary 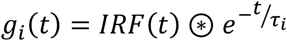, where ⊛ denotes convolution and *τ*_*i*_ is the lifetime of the *i*-th species.

The lifetime values used in the temporal dictionary were estimated directly from the measured time-resolved sinogram through a two-step procedure. First, each radial-angular pixel of the sinogram with sufficient photon counts was fit to a single-exponential decay model after deconvolution with the measured IRF, yielding a per-pixel peak lifetime estimate. Second, the histogram of these per-pixel lifetime estimates was constructed across all sinogram pixels and revealed *M* well-separated modes corresponding to the *M* fluorescent species; the mode locations were taken as the dictionary lifetime values. For the BPAE sample, the histogram exhibited two clear peaks at 1.85 ns and 2.95 ns, attributable to mitochondria and F-actin, respectively. These lifetimes are consistent with values reported in previous studies^31,32^. The estimated lifetimes were then held fixed during the subsequent k-space optimization.

#### Image co-registration

Coarse alignment between the FLIM arm and the reference camera arm was performed during system assembly. Fine spatial registration was achieved in software by estimating an affine transform that maps the reference camera coordinates to the 2D reconstruction grid (226 × 226) used for the FLIM channel, using a fluorescent calibration target with localizable features. The reference image, originally acquired at 1920 × 1080 pixels, was subsequently warped using this affine transform to match the FOV and spatial sampling of the FLIM reconstruction. This procedure ensures that the reference image and the FLIM reconstructions are defined on a common spatial grid for downstream optimization.

### Reconstruction algorithm

#### Forward model

Let *ρ*_*i*_ (*x,y*) denote the spatial concentration map of the *i-th* fluorescent species and *τ*_*i*_ its lifetime. The time-integrated component sinogram is the Radon transform

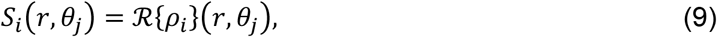

and the measured time-resolved sinogram is modeled as

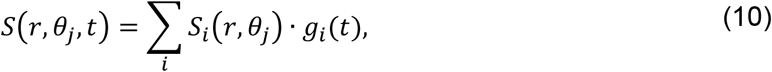

where *g*_*i*_ (*t*) is the IRF-convolved exponential decay kernel for the *i-th* species. The reconstruction goal is to recover {*ρ*_*i*_} from the measured sinogram *S*(*r,θ*_*j*_ *t*), given knowledge of the temporal dictionary {*g*_*i*_} and the high-resolution reference image.

#### Step 1: Per-k temporal unmixing in polar k-space

Let *S* (*r,θ*_*j*,_*t*) denotes the measured time-resolved sinogram (Fig.2a). At each spatial location (*r θ*_*j*_) we solve a regularized linear least-squares problem along the time axis to recover the per-component coefficients *a* ∈ ℝ^*M*^:

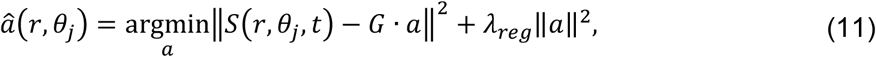

where *G* is the temporal dictionary matrix whose columns are the L^2^-normalized vectors {*g*_*i*_ (*t*) }, and *λ*_*reg*_ is a small Tikhonov regularization parameter that stabilizes the inversion at low-photon spatial coordinates. Because both *S*(*r,θ*_*j*,_*t*) and G are real-valued, the recovered coefficients *â*_*i*_(*r,θ*_*j*_) are also real-valued and represent the photon contribution of component *i* at that spatial location. Any negative entries arise from measurement noise are clipped to zero to enforce strict non-negativity of photon counts, yielding the per-component sinograms *S*_*i*_(*r,θ*_*j*_) at the measured 90 projection angles (Fig.2c,e).

Each per-component sinogram is then transformed into its k-space representation by a 1D Fourier transform along the radial direction r:

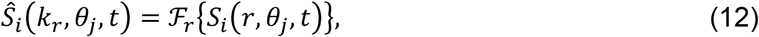

where *ℱ*_*r*_{·} denotes the 1D Fourier transform along r, and *k*_*r*_ is the corresponding radial spatial frequency. By the Fourier slice theorem, *Ŝ*_*i*_ (*k*_*r*_,*θ*_*j*_,*t*) corresponds to one slice of the 2D Fourier transform of the underlying image at angle *θ*_*j*_, time-resolved at *t* (Fig.2 d,f). These complex k-space slices serve as the input to the joint optimization in Step 3.

#### Step 2: Reference spectral prior

The affine-registered reference image *I*_*ref*_ (*x,y*) is Fourier-transformed to obtain its 2D power spectrum:

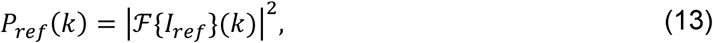

where *k* = (*k*_*x*,_*k*_*y*_) is the 2D spatial frequency vector and *ℱ*{·} denotes the 2D Fourier transform. The power spectrum is normalized so that max *P*_*ref*_ = 1. The spectral weight is then defined in Eq. (1), where *ε* is a small positive constant that prevents division by zero when *P*_*ref*_ is near zero, and *γ* ∈ (0 1] controls the steepness of the weight as a function of reference power. Both values are reported in Supplementary Table S1. The weight *w(k)* is small at frequencies where the reference carries significant structural energy (these frequencies are preserved during reconstruction) and large at noise-dominated frequencies (these are suppressed). Because *w(k*) is defined on a Cartesian k-space grid and the optimization in Step 3 is carried out on a polar grid, we resample *w(k*) onto the polar coordinates (*k*_*r*,_*θ*_*j*_) by bilinear interpolation, yielding *w(r, j)*.

#### Step 3: Polar k-space optimization

The angular sampling is densified from the 90 measured angles to a denser angular grid by linear interpolation in *θ*; in this work the densification factor was eight (720 angles), which we found to balance angular resolution with computational cost. We verified that further densification did not measurably alter the reconstructed images. Let *A(r, j)* denote the total k-space coefficients (the sum across all components) on the dense grid, with *r* indexing the radial spatial frequency and *j* indexing the densified angle. Let *Â*(*r,j*) denote the measured values from Step 1, defined only at the original measurement angles indexed by Ω ⊂ {1 ⋯ 720}. We minimize:

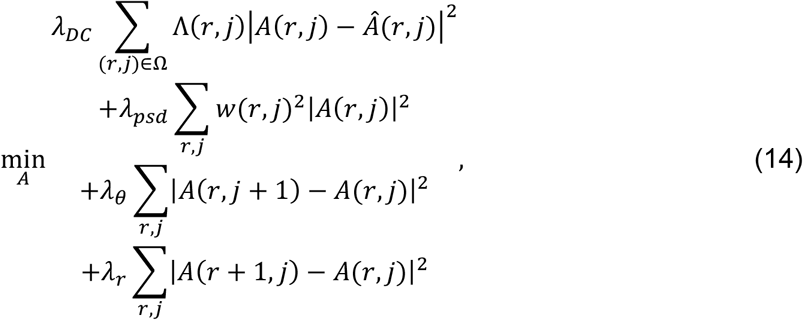

where Λ (*r,j*) is a per-k confidence weight derived from the local signal energy of the measured Fourier slice, and *λ*_*DC*_, *λ*_*psd*_, *λ*_*θ*_, *λ*_*r*_ are scalar regularization coefficients (Supplementary Table S1). The four terms enforce, respectively: (i) data fidelity at the measured angles, (ii) spectral consistency with the reference prior, (iii) angular smoothness across the densified grid, and (iv) radial smoothness. The objective is solved by gradient descent with a fixed step size for a fixed number of iterations, with a non-negativity projection applied periodically (negative values in the inverse Radon transform of *A* are clipped to zero) to enforce physical realizability of the underlying image. The resulting optimized total k-space coefficients are denoted *A*(r, j)*.

#### Step 4: Image-domain fraction and component reconstruction

The optimization in Step 3 is applied only to the total k-space coefficients. To recover individual component images, the per-component sinograms *S*_*i*_ (*r,θ*_*j*_) from Step 1 are first inverse-Radon-transformed (with Ram-Lak filter) at the original 90 measured angles to yield per-component measurement images 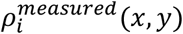. Image-domain fraction maps *f*_*i*_ (*x,y*) are then computed from the per-component measurement images according to Eq. (7). The optimized total k-space coefficients *A*^*^(*r, j*) from Step 3 are inverse-Fourier-transformed along *k*_*r*_ and then inverse-Radon-transformed (Ram-Lak filter) to yield the optimized total image 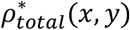. The final component images are then obtained by image-domain multiplication according to Eq. (8). The inscribed-circle output of the inverse Radon transform yields a 226 × 226 reconstructed image at the unchanged sample-plane pixel size (1.74 µm for the beads, 0.87 µm for BPAE). This image-domain fraction approach ensures that the spatial localization of each species is determined entirely by the SPAD measurements through the temporal unmixing of Step 1, while the reference image contributes only to suppressing sparse-view artifacts in the total intensity image through the polar k-space optimization of Step 3.

#### FBP comparison baseline

All FBP reconstructions reported in this work used the same per-k temporal unmixing (Step 1) as the *k*-space fusion method to recover per-component sinograms; the resulting component sinograms at the original 90 angles were then directly inverted by filtered back-projection with a Ram-Lak filter, without any of the spectral-prior optimization steps. This isolates the contribution of the reference-guided polar k-space optimization while keeping the temporal-unmixing step common to both reconstructions, ensuring a fair comparison.

#### Implementation

The reconstruction was implemented in MATLAB R2024a (MathWorks). All Fourier transforms used the FFT routines built into MATLAB. The Step 3 optimization was solved by gradient descent with a fixed step size for a fixed number of iterations; the regularization coefficients *λ*_*DC*_, *λ*_*psd*_, *λ*_*θ*_, *λ*_*r*_ were tuned once on a subset of the data and held fixed across all experiments reported in this work. All hyperparameter values are listed in Supplementary Table S1.

### Sample preparation

#### Fluorescent beads

The samples were prepared from two species of polystyrene microspheres of 15 µm nominal diameter (FluoSpheres, Invitrogen): yellow-green beads with peak excitation/emission at 505/515 nm (catalog F8844) and red beads with peak excitation/emission at 580/605 nm (catalog F8842). Beads of the two species were mixed at approximately equal concentration, sparsely deposited onto a glass coverslip, and allowed to air-dry.

#### BPAE cells

Bovine pulmonary artery endothelial cells (BPAE) were imaged from a commercially prepared slide (FluoCells Prepared Slide #1, Invitrogen). The slide contains fixed and permeabilized BPAE cells stained with three fluorophores; in this work we imaged the two species excitable by our 488/561 nm dual-band illumination: Alexa Fluor 488 phalloidin labeling F-actin filaments and MitoTracker Red CMXRos labeling mitochondria. The third stain (DAPI) was not excited because the excitation path did not include UV wavelengths, ensuring that both the reference image and the FLIM measurement record only the two FLIM-relevant fluorophores. The two fluorophores have distinct fluorescence lifetimes (1.85 ns and 2.95 ns, respectively, in the dictionary used for reconstruction) and are localized to different subcellular structures, enabling simultaneous imaging through the multi-band excitation and emission filters.

### Quantitative evaluation

#### Image quality metrics

##### Signal-to-noise ratio (SNR)

The signal region is defined as the set of pixels where the affine-registered reference image exceeded 30% of its maximum, and the background region is defined as the set of pixels where the reference was below 5% of its maximum. SNR is computed as the ratio of the mean intensity in the signal region of the reconstructed total intensity image to the standard deviation of the reconstructed total intensity image in the background region.

##### Structural similarity (SSIM)

SSIM is computed between the reconstructed total intensity image and the affine-registered reference image, both normalized to unit maximum, using the default MATLAB SSIM function. SSIM measures structural fidelity beyond pixelwise intensity differences and ranges from 0 (no similarity) to 1 (identical images).

##### Contrast-to-noise ratio (CNR)

CNR is computed per component. For component *i*, the signal mask is defined as the set of pixels where the *k*-space fusion-method reconstruction *ρ*_i_ exceeded 10% of its maximum, and the background mask is defined as the set of pixels where the sum *ρ*_1_ + *ρ*_2_ was below 2% of its maximum. CNR for component *i* is then

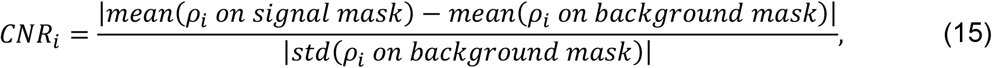

and the reported CNR is the average over the *M* components. The same masks, derived from the *k*-space fusion-method reconstruction, are applied to both the FBP and the k-space fusion-method reconstructions to ensure a consistent comparison.

##### Background noise

The background noise level is reported as the standard deviation of the reconstructed total intensity image in the background mask defined above. Lower values indicate cleaner reconstruction.

## Data availability

The data that support the findings of this study are available from the corresponding author upon reasonable request.

## Use of AI tools

An AI-based language model was used solely to assist with English language editing. All revisions were carefully reviewed by the authors to ensure that the scientific meaning was not altered.

## Ethics declarations

The authors declare that this study does not involve any human participants, animals, or personal data.

