## Supplementary for "High-resolution image-projection fluorescence lifetime imaging microscopy"

### Supplementary Information

#### S1. Numerical simulation of operating regime

To map the operating regime of the proposed reconstruction beyond the two-component biological samples shown in the main text, we performed numerical simulations using synthetic phantoms with controlled lifetime separation $\Delta\tau$ and number of species $M$. The simulations used the same forward model and reconstruction pipeline as the experimental data. Specifically, a sparse set of 90 angular projections was generated from a synthetic bead phantom, convolved with the measured instrument response function and assigned exponential decay kernels, corrupted with Poisson photon-counting noise, and reconstructed using the proposed three-step pipeline. The same hyperparameters used for the experimental data (Section S2) were applied without retuning. For each condition, we report three diagnostics: the conditioning of the temporal dictionary, cond($G^TG$), the normalized lifetime root-mean-square error τ-RMSE / $\Delta\tau$, and the per-bead component contrast ratio defined in Methods. Together, these metrics capture the difficulty of temporal unmixing, the accuracy of the recovered lifetime values, and the spatial fidelity of per-component assignment.

For simulation studies in which the true species at each bead location is known, we additionally computed a per-bead contrast ratio to quantify the specificity of component assignment. For each bead centered at pixel $(y_b, x_b)$ with ground-truth species index $i^{true}$, the contrast ratio is defined as

$$contrast_b = \frac{a_{i^{true}}(y_b, x_b)}{\sum_{i=1}^{M} a_i(y_b, x_b)}, \tag{S1}$$

where $a_i(y, x)$ is the reconstructed amplitude of the $i$-th component at pixel ($y$, $x$), and $i^{true} \in \{1, \cdots, M\}$ denotes the ground-truth species index. In practice, amplitudes were integrated over a ±2-pixel window centered at each bead to account for sub-pixel localization. The overall contrast ratio was then computed as the average across all $N_b$ beads:

$$contrast = \frac{1}{N_b} \sum_{b=1}^{N_b} contrast_b. \tag{S2}$$

The contrast ratio is bounded between 0 and 1. A value of 1 indicates perfect species assignment, with all reconstructed amplitude at each bead assigned to the correct species. A value of 1/$M$ indicates a uniform distribution across the $M$ components, corresponding to random assignment. Because this metric is independent of absolute amplitude scale, it is well suited for comparing component-separation performance across simulation conditions with different lifetime separations or different numbers of species.

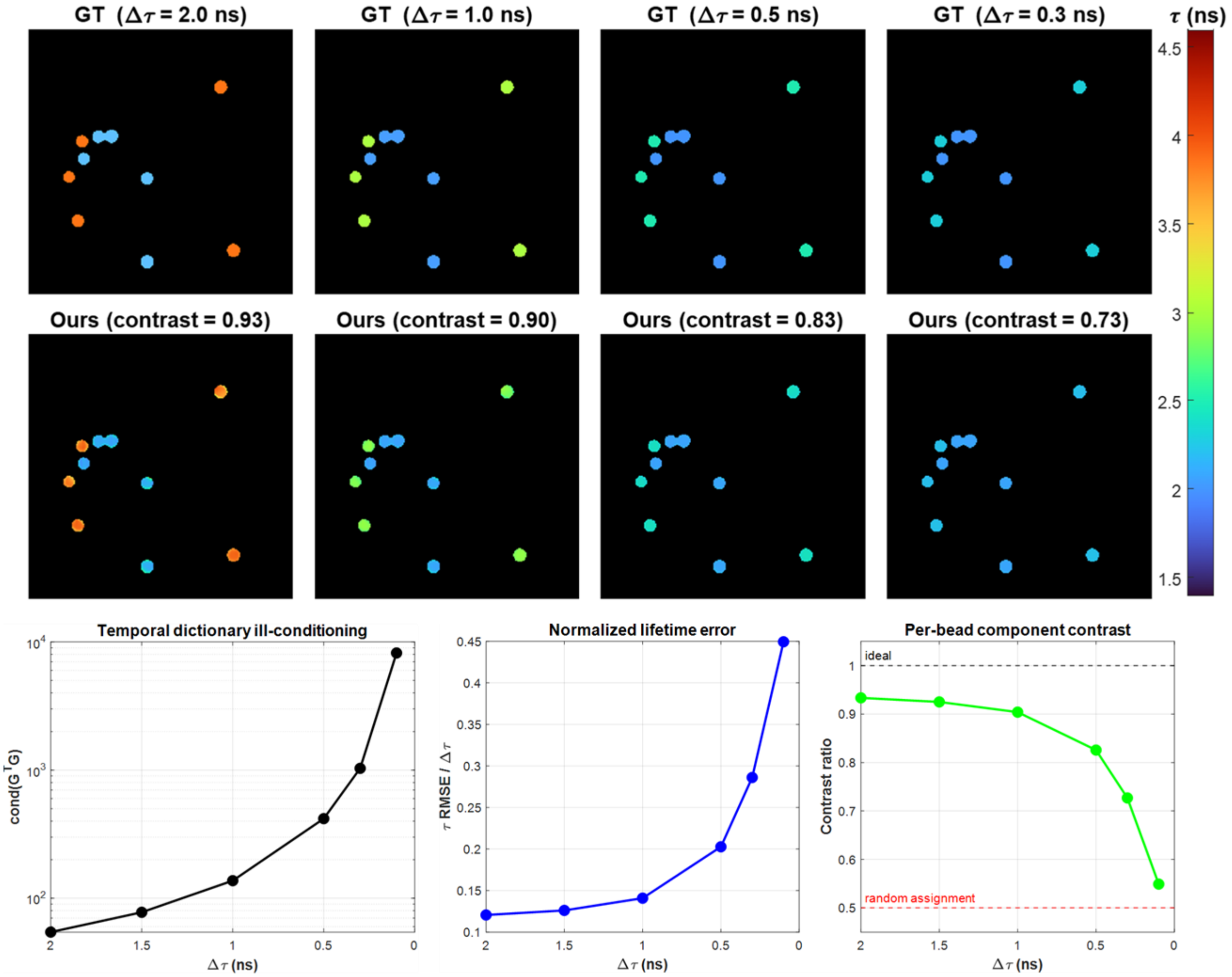


**Figure S1. Effect of lifetime separation $\Delta\tau$ on reconstruction quality ($M = 2$).** Two-component synthetic bead phantoms were reconstructed with the proposed method at seven values of $\Delta\tau \in \{2.0, 1.5, 1.0, 0.7, 0.5, 0.3, 0.1\}$ ns, with all other parameters held fixed. **Top row:** ground-truth lifetime maps for $\Delta\tau$ = 2.0, 1.0, 0.5, and 0.3 ns. **Middle row:** reconstructions by the proposed method, with the per-bead component contrast indicated above each panel. **Bottom row, left to right:** conditioning of the temporal dictionary, cond($G^TG$), which increases by approximately two orders of magnitude as $\Delta\tau$ decreases from 2.0 to 0.1 ns; normalized lifetime error, $\tau$-RMSE / $\Delta\tau$, which remains below 0.15 for $\Delta\tau \geq 1.0$ ns and increases sharply below 0.5 ns; and the per-bead component contrast ratio, which decreases from 0.93 at $\Delta\tau$ = 2.0 ns to 0.55 at $\Delta\tau$ = 0.1 ns. Dashed lines indicate ideal separation (1.0) and random species assignment (0.5). The contrast ratio remains above the random-assignment baseline across the entire tested range.

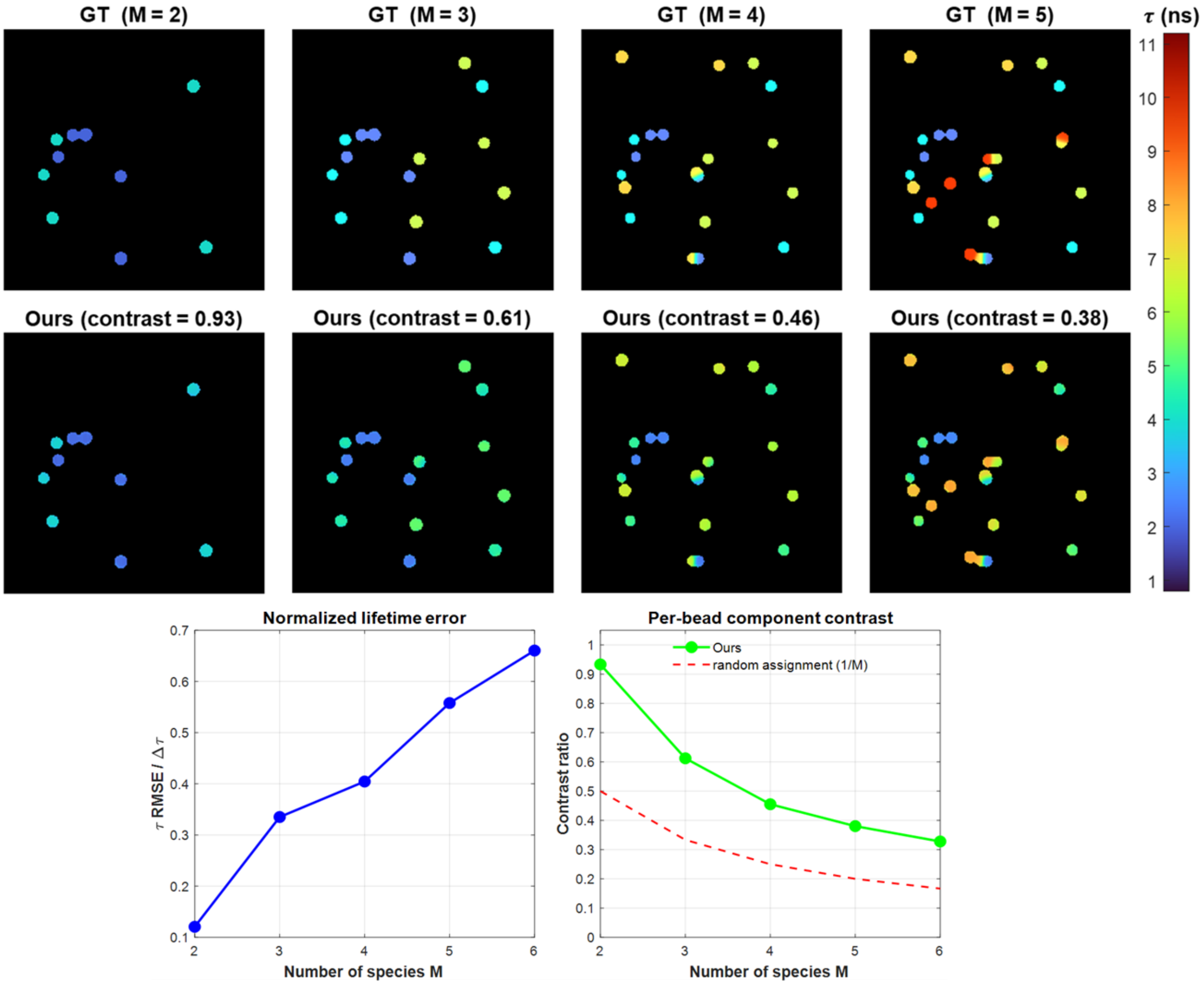


**Figure S2. Effect of the number of species *M* on reconstruction quality ($\Delta\tau$ = 2.0 ns).** Synthetic phantoms containing $M \in \{2, 3, 4, 5, 6\}$ species with lifetimes uniformly spaced by $\Delta\tau$ = 2.0 ns were reconstructed with the proposed method. **Top row:** ground-truth lifetime maps for $M$ = 2, 3, 4, and 5. **Middle row:** reconstructions by the proposed method, with the per-bead component contrast indicated above each panel. **Bottom row:** normalized lifetime error $\tau$ -RMSE / $\Delta\tau$ (left), and per-bead component contrast ratio (right) as functions of *M*. Across all tested values, the contrast ratio remains substantially above the random-assignment baseline of 1/*M*, confirming that the proposed method provides useful component separation even as the number of species increases. The lifetime error increases approximately linearly with *M* because the columns of the temporal dictionary become increasingly correlated as more species are packed into the same temporal range.

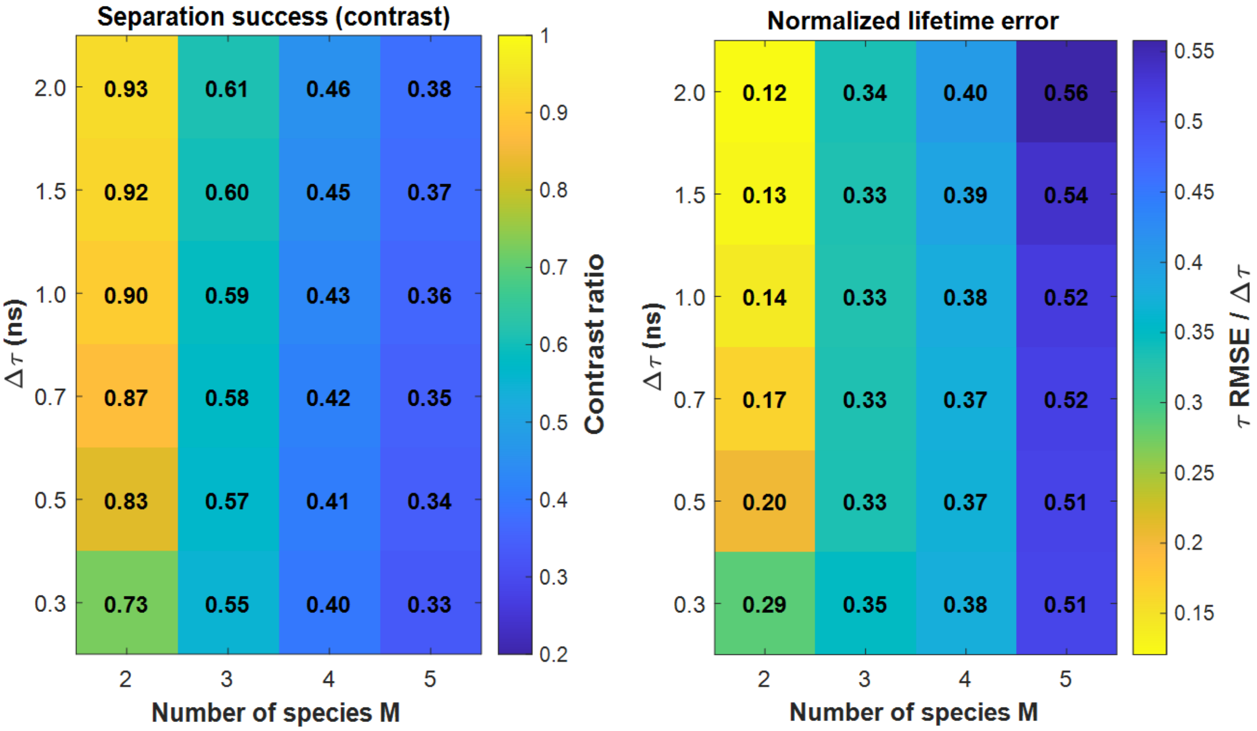


**Figure S3. Two-dimensional operating regime as a function of ($\Delta\tau$, *M*).** The proposed reconstruction was evaluated on a 6 × 4 grid of conditions spanning $\Delta\tau \in \{2.0, 1.5, 1.0, 0.7, 0.5, 0.3\}$ ns and $M \in \{2, 3, 4, 5\}$. **Left:** per-bead component contrast ratio. The contrast ratio is dominated by *M* rather than by $\Delta\tau$: across each column (fixed *M*), the contrast varies only modestly with $\Delta\tau$, whereas across each row (fixed $\Delta\tau$), the contrast decreases by approximately a factor of two as *M* increases from 2 to 5. **Right:** normalized lifetime error $\tau$ -RMSE / $\Delta\tau$ on the same grid. The lifetime error increases with both decreasing $\Delta\tau$ and increasing *M*, reflecting the combined effects of dictionary ill-conditioning and inter-species overlap. Together, these maps define the operating regime in which the proposed method is reliable: the contrast ratio remains above 0.5 for all tested conditions with $M \leq 3$ and degrades gradually outside this regime rather than failing abruptly.

### S2. Hyperparameters and implementation details

Table S1 lists the hyperparameter values used for the proposed reconstruction in all main-text and supplementary results. The values were tuned once on a held-out subset of the bead phantom dataset and then held fixed across both the bead and BPAE experiments, as well as across all simulation conditions in Section S1. All hyperparameters are dimensionless unless otherwise noted.

**Table S1.** Hyperparameters of the proposed reconstruction (BPAE).

| Symbol | Description | Value | Step |
|---|---|---|---|
| $\lambda_{reg}$ | Tikhonov regularization for per-slice temporal unmixing | 0.002 | 1 |
| $\varepsilon$ | Floor in the spectral weight denominator | 0.01 | 2 |
| $\gamma$ | Spectral weight exponent applied to $P_{ref}$(k) | 0.5 | 2 |
| $\lambda_{DC}$ | Data-fidelity weight at measured angles | 8.0 | 3 |
| $\lambda_{PSD}$ | Spectral prior weight | 0.003 | 3 |
| $\lambda_{\theta}$ | Angular smoothness weight | 0.012 | 3 |
| $\lambda_{r}$ | Radial smoothness weight | 0.002 | 3 |
| $N_{\theta,dense}$ | Densified angular grid size | 720 | 3 |
| $N_{iter}$ | Gradient-descent iterations in Step 3 | 700 | 3 |
| $\eta$ | Gradient-descent step size in Step 3 | 0.08 | 3 |

#### Sensitivity to hyperparameters

The reconstruction was most sensitive to the spectral prior weight $\lambda_{psd}$ and the spectral weight exponent γ. Small $\lambda_{psd}$ values reduce the influence of the reference image, producing reconstructions close to the FBP baseline, whereas large $\lambda_{psd}$ values over-smooth high-frequency content that is weak or absent in the reference image. The exponent $\gamma$ controls the steepness of the transition between preserved and suppressed frequencies in *w*(*k*); values near 0 produce nearly uniform suppression, whereas values approaching 1 sharpen the transition and can introduce ringing near edges. The angular and radial smoothness terms $\lambda_{\theta}$ and $\lambda_{r}$, mainly affect noise levels in the optimized sinogram and are relatively insensitive to sample type when set on the same order of magnitude as $\lambda_{psd}$. The data-fidelity weight $\lambda_{DC}$ was fixed at 8.0, and all other weights were expressed relative to this value.

### Computational cost

The reconstruction was implemented in MATLAB R2024a (MathWorks). All Fourier transforms used MATLAB's built-in FFT routines, and no GPU acceleration was used. Reconstructions were performed on a workstation with an Intel Core i7 multi-core CPU and 32 GB of RAM. The dominant computational cost was Step 3 polar k-space optimization, which scales as $O(N_{iter} \cdot N_r \cdot N_{\theta,dense})$, where $N_{iter}$ is the number of gradient-descent iterations, $N_r$ is the number of radial samples, and $N_{\theta,dense}$ is the size of the densified angular grid. For the experimental dataset (320 radial samples, 720 angles, and 700 iterations), the complete four-step reconstruction pipeline completed in approximately 10 s per dataset; Step 3 accounted for 7.4 s. The per-slice temporal unmixing in Step 1 was parallelized over polar coordinates and accounted for a small fraction of the total runtime (0.7 s). The regularization coefficients $\lambda_{DC}$, $\lambda_{psd}$, $\lambda_{\theta}$, and $\lambda_r$ were tuned once on a subset of the bead-phantom data and held fixed across all experiments and simulation conditions reported in this work. All hyperparameter values are listed in Table S1.
